# Decreasing photosystem antenna size: a double-edged sword for canopy photo-synthetic efficiency

**DOI:** 10.1101/2022.06.09.495562

**Authors:** Linxiong Mao, Qingfeng Song, Ming Li, Xinyu Liu, Zai Shi, Faming Chen, Gen-yun Chen, Xin-Guang Zhu

## Abstract

Optimization of antenna size of photosynthetic systems is one strategy to increase plant canopy photosynthesis and crop yield potential. The relationship between antenna size and photosynthesis rate has been extensively studied recently. However, conflicting results have been obtained. Here we show that the extent of decrease in antenna size is a major factor determining the consequences of decreasing antenna on photosynthesis and growth-related parameters. Specifically, we constructed transgenic rice lines with artificial microRNA (amiRNA) targeting to Chlorophyll Synthesis (*YGL1*) to generate transgene heterozygous and homozygous lines with different leaf chlorophyll contents and antenna sizes. We found that canopy photosynthesis (A_c_), biomass and grain yield of the heterozygote were not significantly different from those of WT while the A_c_, biomass and grain yield of the homozygote were lower than those of WT. Further, when the maximal quantum yield of photosystem II (F_v_/F_m_) was larger than 0.8, decreasing antenna size by reducing chlorophyll biosynthesis did not affect leaf photosynthesis; but when F_v_/F_m_ was lower than 0.8, there is a positive relationship between antenna size and leaf photosynthesis. There is large variation in both leaf chlorophyll content and antenna size in elite rice cultivars, suggesting that there is a large scope to decrease leaf chlorophyll content to increase nitrogen use efficiency as long as the quantum yield of PSII is not compromised.

## INTRODUCTION

Crop yield potential is determined by light interception coefficient, energy conversion efficiency and harvest index (Monteith & Moss, 1977). For current high yield crops, the light interception coefficient and harvest index are close to their upper bound, but the energy conversion efficiency is lower than the theoretical maximal and can be improved to increase crop yield potential (Long et al., 2015, Dann & Leister, 2017, Zhu et al., 2010). Approaches of increasing energy conversion efficiency have been explored in major crops, including accelerating recovery from photo-protected state, altering stomatal density, enhancing photorespiratory carbon flow, optimizing light harvesting and *etc*, see review by (Bailey-Serres et al., 2019). The size of antenna, *i*.*e*. the number of chlorophyll (Chl) molecules associated with light-harvesting protein complex (Fromme et al., 2003), influences the efficiency of energy transfer from antenna Chl to reaction center and utilization for photochemistry and hence is recognized as a target for optimization (Melis, 2009). Compared to Chl a, the relative content of Chl b is lower in photosystem reaction center and higher in light harvesting complex, so the Chl a/b ratio is commonly used as an indicator to antenna size (Liu et al., 2004, Liu & Chang, 2008). Based on the biochemical basis, the antenna size can be quantified by the ratio of the amounts of LHCII to D1/D2 (Jansson, 1999, Rochaix, 2014), or by the Chl *a/b* ratio (Tanaka et al., 2001, Masuda et al., 2003). Larger antenna can intercept more photons under the same level of photosynthetic photon flux density (PPFD), resulting in Q_A_ reduction faster and the chlorophyll fluorescence increase faster (Kramer et al., 2004). So, the increasing slope for the fast phase of the Chl fluorescence induction curve, *i*.*e*. the OJIP curve, correlates with antenna size (Preiss & Thornber, 1995, Zhu et al., 2005). Specifically, the fluorescence intensity at 300µs after excitation under different photosynthetic photon flux density (PPFD) presents the functional antenna size (Bielczynski et al., 2020).

There are a number of reported options to manipulate leaf chlorophyll content and antenna size. Antenna size can be decreased when chlorophyll content is decreased. In many of these reported studies, their impact on chlorophyll contents and photosynthetic rates were reported, however, their impacts on antenna sizes are mostly not reported. For example, decreased enzyme activity of chlorophyll synthase (YGL1)(Wu et al., 2007) results in less chlorophyll content and higher Chl *a/b* ratio, which indicates an smaller antenna size, and the mutant *ygl1* with higher leaf photosynthesis rate (Li et al., 2013). Magnesium (Mg)-chelatase catalyzes the transformation of chlorophyll precursors and mutation of *Ygl7* encoding the subunit D of Mg-chelatase can decrease chlorophyll content while increase leaf photosynthesis (Deng et al., 2014). The gene of *ChlH*, which encodes Mg chelatase H subunit, is regulated by GUN4 and affects the content of magnesium protoporphyrin IX (MgP(IX))(Davison et al., 2005) and also chlorophyll content eventually. In soybean, mutation of the gene (*Y11y11*) encoding magnesium chelatase subunit-I (CHLI) (Campbell et al., 2014) results in lower chlorophyll content and similar leaf photosynthesis rate with the WT (Slattery et al., 2017). The photosynthesis rate and quantum yield of photosystem II (Φ_PSII_) of the *Y11y11* are higher and NPQ of the *Y11y11* is lower compared to WT (Slattery et al., 2017, Sakowska et al., 2018). The *CAO* gene encodes chlorophyllide a oxygenase, which is associated with the Chl b synthesis (Morita et al., 2005), and the expression of *CAO* can regulate the antenna size (Tanaka et al., 2001, Masuda et al., 2003). Besides, antenna size is also influenced by the chloroplast signal recognition particle (CpSRP) pathway(Kirst & Melis, 2014).

Altering leaf chlorophyll content and antenna size has been reported to improve photosynthesis. The photosynthesis rate of rice pale green leaf (*pgl*) mutant is higher than WT under high light condition (Gu et al., 2017a, Gu et al., 2017b). At the canopy scale, decreasing leaf Chl content may increase leaf transmittance, improve the light distribution in a plant canopy and improve canopy photosynthesis according to modeling study (Song et al., 2017). Similarly, for algae, the light can penetrate deeper in the culture solution of the smaller antenna mutant *tla1*, resulting in an increased photosynthesis rate per volume (Polle et al., 2003). Moreover, downregulation of *CpSRP43*, a CpSRP pathway gene through RNAi technology in tobacco reduces the Chl content and enhances the biomass production (Kirst et al., 2018).

In contrast, there are also examples showing that photosynthesis rate is decreased in less Chl mutants. For instance, mutation of *OsClpP6* results in impaired photosynthesis apparatus and chloroplast development (Dong et al., 2013). Loss of the transcription factor *OsPIL1* results in decreased chlorophyll content with simultaneous decrease in panicle length, the number of grains per panicle, and thousand grain weight comparing to WT (Sakuraba et al., 2017). These results show that decreasing antenna size might either promote or decrease photosynthesis.

In this study, we constructed transgenic rice lines with different Chl content and antenna size with artificial microRNA (amiRNA) targeting to Chlorophyll synthesis (*YGL1*), then we studied the impacts of altering antenna size on leaf and canopy photosynthesis, biomass and grain yield. We test the hypothesis that these different responses of decreasing antenna size on leaf photosynthesis are determined by extent of decrease in antenna size, *i*.*e*. only when the antenna size is in certain range, decreasing antenna size can promote photosynthesis.

## MATERIALS AND METHODS

### Construction of YGL1-RNAi suppression vector

To construct the RNAi vector of Chlorophyll synthesis (*YGL1*, Os05g0349700), Web MicroRNA Designer platform (WMD) (http://wmd3.weigelworld.org/cgi-bin/webapp.cgi) was used to design amiRNA (Artificial microRNA) sequence (21 mer) with reference genome “Oryza sativa cDNA v5 (TIGR)”. The amiRNA sequence TTTATCGCATCTATATCTCGG, designed with WMD, was used to construct a vector. The amiRNA precursor was amplified by overlapping PCR from the pNW55 template (containing the osa-Mir528 precursor) to replace the 21 bases of the natural osa-MIR528 miRNA as well as the partially complementary region of the miRNA in a similar way as described in previous study (Schwab et al., 2006). First, three fragments including the multiple cloning sites (MCS) were amplified by PCR from the template clone pNW55 using three pairs of primers (Table S1), (G4368, Primer II), (Primer I, Primer IV) and (Primer III, G4369). Second, the three fragments were fused by one fusing PCR with the two flanking primers G4368 and G4369. All PCRs were performed with Planta Super-Fidelity DNA polymerase (P505, Vazyme, China) and PCR primers were shown in Table S1.

### Plant materials

Rice (*Oryza sativa* spp *japonica* cv Nipponbare) was transformed by the agrobacterium mediated methods following (Hiei et al., 1994). Insertion sites in chromosome were identified by TAIL-PCR (Tan et al., 2019) with universal primers mLAD1, mLAD2, mLAD3, mLAD4, AC0 and AC1 and specific primers TAIL-LB-1, TAIL-LB-2 and TAIL-LB-3 (Table S1). Heterozygous and homozygous plants of two transgenic lines (line 7 and 8) were identified by PCR with primers (Y7-genome-F-3, Y7-genome-R-3, Y7-genome-R and TAIL-LB-1) for line 7 and (Y8-genome-2R, Y8-zaiti-F and Y8-genome-F) for line 8 (Table S1). Heterozygous and homozygous plants of the two transgenic lines were used to conduct pot experiments and field trails.

Rice high yield cultivars HuangHuaZhan (HHZ), 9311, HuangGuangYouZhan (HGYZ), HuangGuangYouZhan 2(HGYZ2), ChaoYou 1000 (CY1000) and Y-LiangYou 2 (YLY2) were used in the field trails.

### Plant growth conditions

The plants were grown in paddy field at the experimental site of the Shanghai Institute of Plant Physiology and Ecology (121°8′1.33″E, 30°56′26.73″N). The rice seedlings were transplanted to paddy soil on June the 21^th^ in 2019 and on June the 30^th^ in 2020. Standard agronomic practices were applied for field management. 400 kg/ha fertilizer (15% N, 15% P_2_O_5_, 15% K_2_O) was applied before transplantation.

In green house, rice were grown in pots under an alternating light/dark cycle (12h light and 12h dark), and light intensity was about 400 μmol m^−2^ s^−1^. The temperature of the green house was set at around 28 °C.

### RNA extraction, RT-PCR/qRT-PCR

Total RNA was extracted from rice leaves at the booting stage using a Trizol reagent (15596018, Thermo Fisher Scientific, USA). RT-PCR was performed with a One-Step RT-PCR kit (AT311-03, Transgen biotech, China) on 50 ng total RNA with random primers and stem-loop primer, which is designed for the amiRNA (Schwab et al., 2006). Primers for qRT-PCR were designed with PrimerQuest Tool IDT (https://sg.idtdna.com) and Real-time PCR was performed using a SYBR Premix Ex TaqTM kit (RR01CM (AM × 12), Takara Bio Inc., Japan) to evaluate the mRNA level of *YGL1* with actin as internal control on Bio-Rad CFX-96. The 20 μl PCR reaction mixture consisted of Taq DNA Polymerase (AQ141-02, Transgen biotech, China), 10× PCR buffer, 1.5 μg of cDNA template, 0.5 μM of each primer. Reaction mixtures (for both *YGL1* and amiRNA) were initially heated at 95°C for 3 min followed by 42 cycles for 10 sec at 95°C, 20 sec at 58 °C, 30 sec at 72 °C, and finally 5 sec at 65 °C and 30 sec at 95 °C. qRT-PCR was carried out three times to ensure reproducibility of the results. The transcripts expression levels of *YGL1* and amiRNA were quantified relatively to those of the wild type, normalized to the expression level of *OsActin1*. The quantification of amiRNA was referred as before (Chen et al., 2005, Varkonyi-Gasic et al., 2007). All the primers used above were shown in Table S1.

### Protein extraction and Western Blot

Total protein of rice leaves at booting stage was extracted and stored with 5×SDS loading buffer. Before western blot, the total protein was extracted and put into 95°C for 5 minutes, then loaded on SDS-PAGE for immunoblotting. The protocol of SDS-PAGE and the reference proteins of immunoblotting followed previous study (Li et al., 2011). Primary antibodies against proteins including antenna (Lhcb1, Lhca3), PSII core subunit D1 (PsbA), PSI core subunit (PsaA), peripheral protein (PsbO) and cytochrome (PetA, PetC). Antibodies are prepared with peptides of these proteins (peptides sequences are shown in Table S2).

The second antibody is Goat Anti-Rabbit IgG H&L (HRP) (ab6721, Abcam). Western blot images were captured with a CCD imager (ChemiDoc Touch, Tanon) with Western Blotting Detection Reagents (Thermo Fisher Scientific, USA). The concentration of soluble protein was determined with the Bradford kit (Dalian Meilun Biological Technology, China).

### Chlorophyll determination

Chlorophyll and other pigments were extracted with 80% acetone according to the protocol (Croce et al., 2002). The content of chlorophyll a and b was determined with spectrophotometer. The SPAD (502 Plus, Konica Minolta, INC. Japan) was also used for measuring leaf chlorophyll content in the field.

### Leaf photosynthesis measurement and data fitting

Leaf photosynthetic gas exchange measurement was performed with the portable leaf gas exchange system LI-6400 (Li-Cor Inc, Lincoln, Nebraska, USA). Leaf photosynthetic CO_2_ assimilation rates for the new fully expended leaves at booting stage (40 DAT, days after transplanting) and flag leaves at grain filling stage (65 DAT) were measured under saturate light conditions (PAR 1800 μmol m^−2^ s^−1^, CO_2_ 400 μmol mol^−1^), the temperature in the leaf chamber block was set to be the same as ambient temperature (36 °C for booting stage, 34 °C for grain filling stage), humidity in the leaf chamber was controlled at around 37% to 42% at booting stage and at around 34% to 42% at grain filling stage. Light response curves of leaf photosynthetic CO_2_ assimilation rate (A-Q curves) were measured with step changing of photosynthetic photon flux density (PPFD) from 1800 to 0 μmol m^−2^ s^−1^ (1800, 1600, 1200, 1000, 800, 600, 400, 300, 200, 150, 100, 50 and 0), the other setting was the same as above. CO_2_ response curves of leaf photosynthetic CO_2_ assimilation rate (A-Ci curves) were measured with step changing of CO_2_ concentration ([CO_2_]) from 425 to 50 μmol mol^−1^ (425, 350, 250, 200, 150, 100, 50), then [CO_2_] was set to be 425 μmol mol^−1^ for 10 minutes and then increase to 1800 μmol mol^−1^ (425, 500, 700, 900, 1100, 1300, 1500, 1800). The plants including root with soil were carefully isolated from paddy field to pots (20 L volume) the day before leaf gas exchange measurement following a protocol (Chang et al., 2017).

Parameters of A-C_i_ curves, V_c,max_ (maximal carboxylation rate under RuBP and CO_2_ saturation, unit: µmol m^−2^ s^−1^), J_max_ (maximal linear electron transport rate under saturate light, unit: µmol m^−2^ s^−1^) were fitted with the FvCB biochemical model (Farquhar et al., 1980).

### Canopy photosynthesis measurement with a multi-chamber canopy gas exchange system

Canopy photosynthesis rate was measured with a canopy photosynthesis and transpiration system (CAPTS) (Song et al., 2016, Song & Zhu, 2018), with the protocol used in previous studies (Chang et al., 2019). The CAPTS system includes one console connecting ten chambers and the chamber size is 1m×1m×1.2m (L × W × H). The chambers are open and closed automatically controlled by the console. The wall of the chamber is made of transparent polycarbonate film (1.5 mm thickness, 75% transmittance). The lid of the chambers is made of glass (5 mm thickness, 85% transmittance) and can be fully opened (the angle of lid changed from horizontal to vertical). There were four fans installed inside of each chamber for gas mixing. In the console, there is a multiplexer for switching gas flow from one of the ten chambers and pump to the infrared gas analyzer (IRGA). The CO_2_ concentration was measured and recorded by the console with an interval of 1s.

During the measurement of canopy gas exchange, plants in the field were enclosed in the chamber. One of the ten chambers was closed for two minutes for a measurement, and after, the chamber was opened, and the next chamber was closed for next measurement. The measurement was performed in a rotating order to minimize the potential impact of measurement sequence on the results, specifically, we start from chamber one to ten and then goes back to the chamber one. The data recorded by the console were used to calculate the rate of CO_2_ concentration change with time (*dc/dt*). The net CO_2_ flux (*F*_*c*_) including canopy, root and soil flux, was calculated with Eqn 1, as used in previous studies (Song et al., 2016)(Steduto et al., 2002).

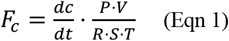

Where *P* (unit: kPa) is air pressure in the canopy chamber, *V* (unit: m^3^) is the volume of air in the chamber, *S* (unit: m^2^) is the ground area covered by the canopy, *T* (unit: K) is the air temperature and *R* (8.314×10^−3^ m^3^ kPa mol^−1^ K^−1^) is universal gas constant.

In this study, we used 20 chambers associated with two consoles, with two chambers measuring paddy soil respiration, and with 18 chambers measuring the flux of whole plant (canopy plus root) and soil together.

### Measurements of agronomic traits

Agronomic traits including plant height, number of panicles per plant, grain number per panicle, above ground biomass dry weight and panicle dry weight were measured at harvest stage. Plant height was the distance from ground surface to the tip of flag leaf. Above ground biomass was harvested and dried at 110 °C for 1h and 70°C for 3 days. The panicles were dried at 37°C for more than 1 week. 28 biological replicates were used for plant height, panicle number per plant and biomass and six biological replicates were used for the weight of panicle.

### Quenching analysis, high light treatment and dark recovery experiment

DUAL-PAM-100 fluorometer (Walz, Germany) was used to measure chlorophyll fluorescence. Chlorophyll fluorescence parameters, such as maximal or potential photochemical efficiency of photosystem II in dark-adapted leaves (F_v_/F_m_), nonphotochemical quenching (NPQ), Φ(PSII), Φ(NPQ), were calculated according to (Kitajima & Butler, 1975, Genty et al., 1989). The equations to calculate these parameters are shown as follows (Eqn 2-6).

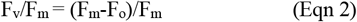

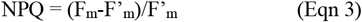

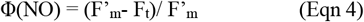

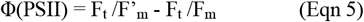

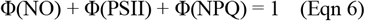

where the F_o_ is minimum fluorescence yield, F_m_ is maximum fluorescence yield, F_v_ is variable fluorescence and is calculated as F_m_-F_o_, F_t_ is steady-state fluorescence yield, F’_m_ is partial open maximum fluorescence yield (Kate Maxwell, 2000).

For the high light treatment and dark recovery experiments, the leaves were adapted under dark for more than two hours and the middle segments of leaves were cut and put into a wet gauze. The PPFD of the high light treatment was 1131μmol m^−2^ s^−1^, F_v_/F_m_ was measured every 1.5 hours of during the light treatment or dark recovery. Before measurement of each F_v_/F_m_, a 5 min dark recovery was applied.

### Functional antenna size of PSII measurement

The PSII functional antenna size was measured with the method used in previous studies (Dinç et al., 2012, Bielczynski et al., 2016) with Pocket-PEA (Hansatech, U.K.). The OJIP curve represents the transient fast chlorophyll a fluorescence induction of the dark-adapted leaf following excitation with 1 s of saturating orange-red light which were described in detail in (Essemine et al., 2020). Plants were dark adapted for more than 1 h before measurements. Then, leaves which were used to measure leaf photosynthesis, were exposed to saturating orange-red (625 nm) under different actinic light (600, 700, 800, 900, 1000, 1200, 1500, 2000, 2500, 3000, 3500μmol m^−2^ s^−1^) provided by the LED for 1 second.

Between these OJIP measurements under different PPFD, the leaf was dark adapted for more than 10 minutes. The slope of linear regression between relative fluorescence signal at 300 μs (F_300µs_/PPFD) and the PPFD was defined as the functional antenna size (Dinç et al., 2012, Bielczynski et al., 2016).

### Statistical analysis

All data in the text, tables, and figures are reported as mean ± standard deviation (sd), unless otherwise indicated. An unpaired Student *t-test* statistical analysis was used to determine the statistical significance between different transgenic lines and wild-type. All statistical analysis were performed with *R* software (version 3.5.3) (https://www.r-project.org/).

## RESULTS

### Leaf chlorophyll content correlates with *YGL1* expression level in transgenic rice

To study the relationship among chlorophyll content, antenna size and photosynthesis, we used the pNW55 vector with 35s promoter to construct transgenic rice lines (Fig. 1a). Five Ami-*YGL1* (artificial miRNA targeting *YGL1*) transgenic rice lines were generated, and two lines were selected for our study. The expression levels of amiRNA in homozygous lines were significantly higher than those in the heterozygous lines (Fig. 1b), correspondingly the expression levels of *YGL1* in homozygous lines were lower than those in the heterozygous lines (Fig. 1c). The expression levels of *YGL1* for both homozygous and heterozygous were significantly lower (*P* < 0.001) than those in the WT (Fig. 1c). The amounts of YGL1 protein determined with western blot for these Ami-*YGL1* lines were consistent with the results of mRNA expression levels (Fig. 1d), confirming that *YGL1* was effectively downregulated by the amiRNA transgene. The chlorophyll content was measured with SPAD (Fig. 1e) and the SPAD value correlates with the expression levels of *YGL1* (Fig. 1f) with R^2^=0.6215. The phenotype of the plants and leaves are shown in Fig. 1G-H, where leaves were pale green for heterozygote, yellow green for homozygote and dark green for WT (Fig. 1g∼h).

**Figure 1.**
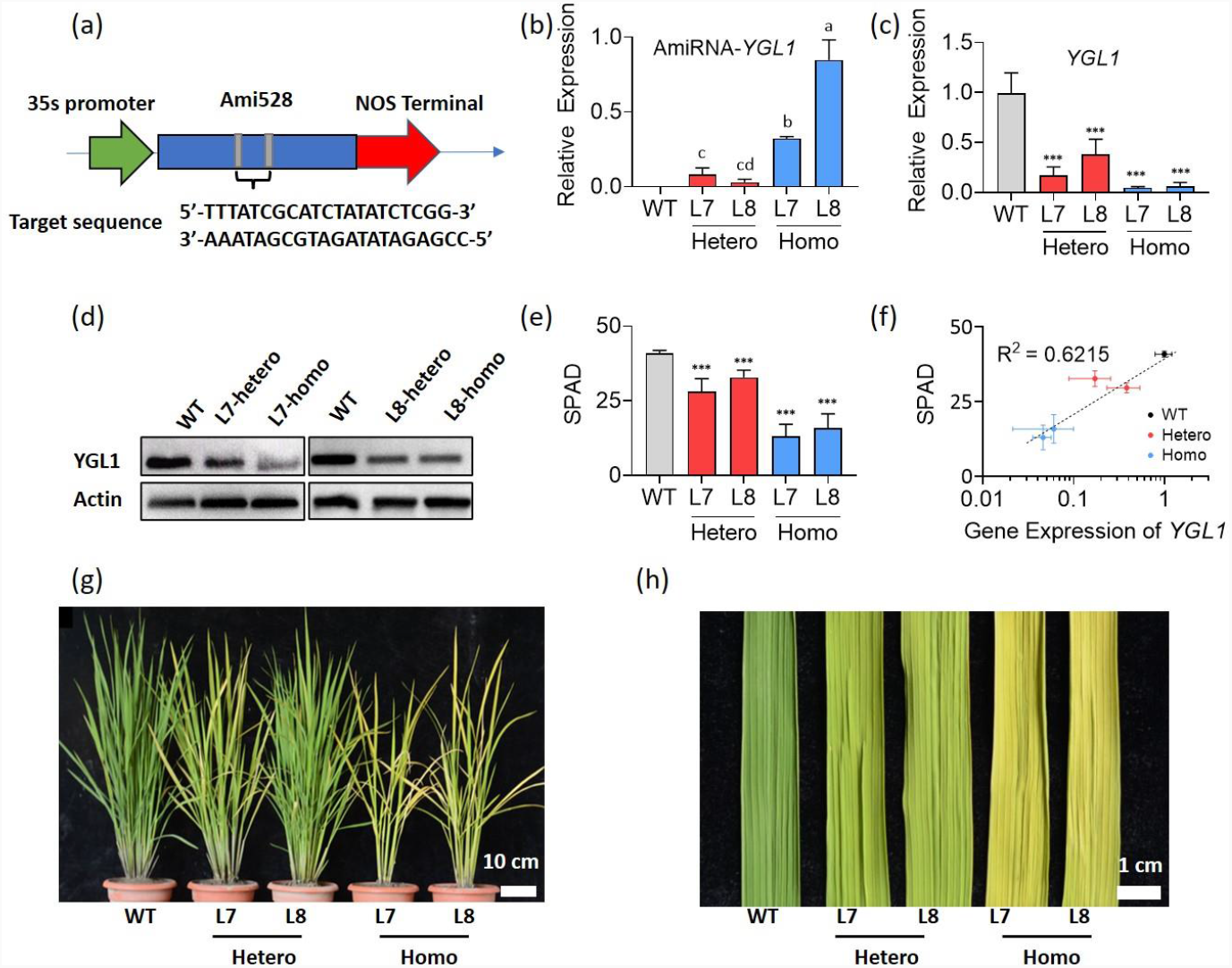
The construction and phenotype of Ami-*YGL1* rice. Schematic presentation of pNW55-RNAi plasmid for the Ami-*YGL1* gene in rice **(a)**. Expression level of amiRNA **(b)** and the targeting gene *YGL1* **(c)** in Ami-*YGL1* heterozygous, homozygous lines and the WT plants. Total RNA was extracted with Trizol from leaves at the tillering stages and expression levels were analyzed with qRT-PCR and *OsActin1* was used as an internal control to normalize the expression levels. Data were shown as mean+/− std (n=3 plants). **(d)** Amounts of YGL1 in rice leaves at the tillering stage for the Ami-*YGL1* homozygous, heterozygous lines and WT was determined with western blot. **(e)**The chlorophyll content in WT, Ami-*YGL1* heterozygous and homozygous lines was represented by SPAD, data were shown in mean +/− std (n = 5 plants) and triple asterisk shows Student’s *t*-test *P*<0.001. **(f)** The correlation between SPAD and the expression of *YGL1* in WT and Ami-*YGL1* lines. **(g)** Phenotype of individual plants and **(h)** the new fully expended leaves from different transgenic lines and WT.

### Antenna size was decreased in the Ami-*YGL1* lines

The changes of antenna size in the Ami-*YGL1* heterozygous and homozygous lines were studied with amounts of proteins detected with western blot, chlorophyll (Chl) a/b ratio and functional antenna size estimates. Chl a/b ratio was increased with decreasing YGL1 (Fig. 2a), indicating that the light harvesting complex (LHC) binding pigments decreased more than the photosystem core (PC) binding pigments because Chl a/b ratio for LHC is lower than PC. Results of immunoblotting analysis further supports that the LHC proteins were decreased more than the PC proteins, and specifically, the amounts of antenna proteins, Lhcb1 and Lhca3, in Ami-*YGL1* lines were less than WT (Fig. 2b). Besides, the amount of photosystem II core subunit D1 kept the same in Ami-*YGL1* heterozygote and decreased in homozygote compared to WT (Fig. 2b). The amounts of photosystem I core subunit PsaA, peripheral protein PsbO and cytochrome subunits PetA and PetC in Ami-*YGL1* lines were the same as WT (Fig. 2b). Further, functional antenna size of PSII was quantified with chlorophyll fluorescence measurement for field grown plants. And calculated from the chlorophyll fluorescence OJIP curves (Supplementary Fig S1) measured following the methods in previous studies (Dinç et al., 2012, Bielczynski et al., 2016). The functional antenna size of heterozygous lines was 75% (line 7, 75%±6%), 76% (line 8, 76%±9%) of WT and homozygous lines was 43% (line 7, 43%±6%), 57% (line 8, 57%±8%) (Fig. 2c).

**Figure 2.**
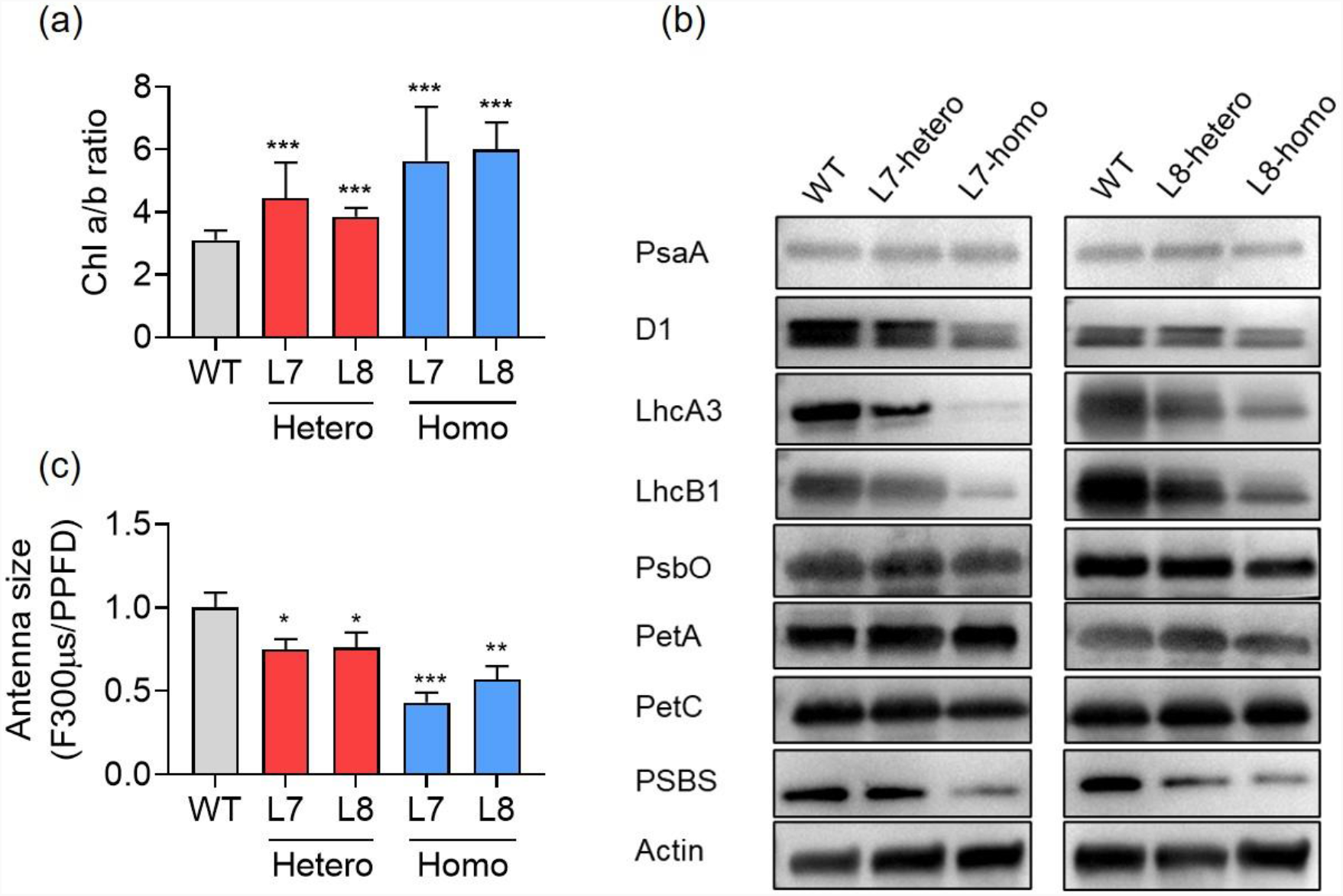
The chlorophyll a/b ratio, amounts of proteins/protein complexes and functional antenna size for Ami-*YGL1* rice lines. **(a)** Chlorophyll a/b ratio. **(b)** Functional antenna size of PSII quantified with the slope of chlorophyll fluorescence at 300 μs against PPFD. Chlorophyll fluorescence was normalized with induction PPFD. **(c)** Amounts of proteins detected with western blot. Data were shown with mean +/− std (n=8-12 plants) and asterisk shows Student’s *t*-test *P*<0.001 (***), *P*<0.01 (**) *P*<0.05 (*).

### Gas exchange and chlorophyll fluorescence measurement of Ami-*YGL1* lines

Light response (*A-Q*) curves and CO_2_ response (*A-C*_*i*_) curves for leaf photosynthesis were measured for the field grown rice plants at both tillering and grain filling stage to assess the effects of changing antenna size on photosynthesis in the Ami-*YGL1* heterozygous and homozygous plants. At tillering stage, leaf photosynthetic CO_2_ assimilation rate (a) was not significantly different between the heterozygote and WT under different PPFD levels and intercellular CO_2_ concentrations (C_i_) at tillering stage (Fig. 3a-b, Supplementary Table S3). *A* for homozygote was lower than WT under high PPFD levels and under *C*_*i*_ higher than about 350 µmol mol^−1^ (Fig. 3a-b, Supplementary Table S3). Maximal Rubisco carboxylation rate under saturate CO_2_ and RuBP (*V*_*cmax*_) and maximal electron transport rate (*J*_*max*_) was fitted from *A-C*_*i*_ curves using FvCB model and the *V*_*cmax*_ and *J*_*max*_ were both lower in line 8 homozygote than other lines (Supplementary Table S4). We further assessed the photochemical efficiency of PSII (Φ_PSII_, Fig. 3c) and nonphotochemical quenching (NPQ, Fig. 3d) between these lines. Our results show that the NPQ was the highest in WT, medium in heterozygote and lowest in homozygote, while the Φ_PSII_ was the lowest in WT, medium in heterozygote and highest in homozygote (Fig. 3c, d).

**Figure 3.**
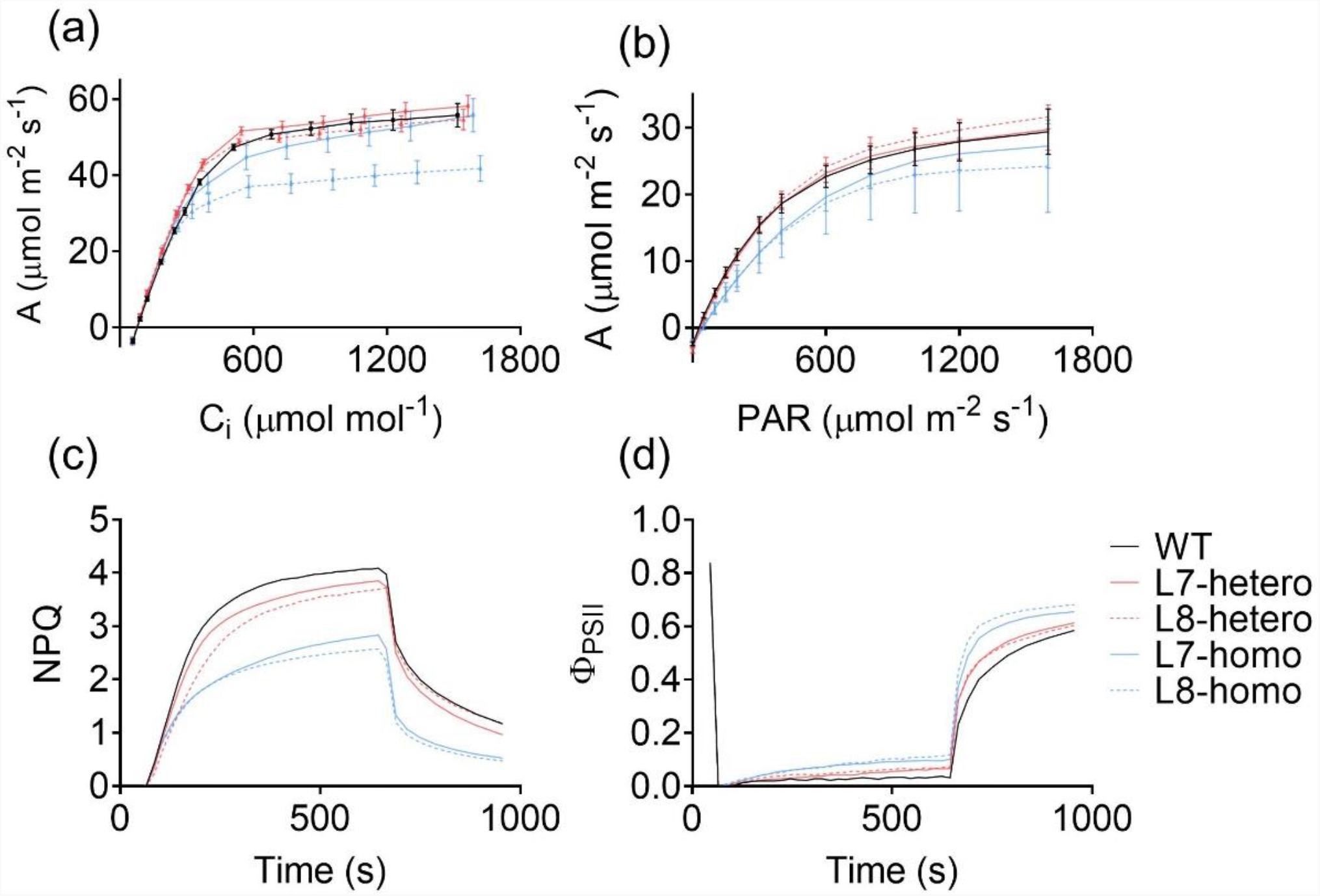
CO_2_ response curve **(a)** and light response curve **(b)** of leaf photosynthesis. The Φ_PSII_ **(c)** and NPQ **(d)** kinetics on Ami-YGL1 lines and WT leaves illuminated with 1,316 μmol photons m^−2^ s^−1^ for 10min followed by 5min dark.

### Canopy photosynthesis and canopy light absorbance of Ami-*YGL1* lines in field

Diurnal curves of canopy photosynthetic CO_2_ uptake rate (*A*_*c*_) (Fig. 4a, d) and whole day cumulative CO_2_ uptake of the whole canopy (*A*_*c,d*_) (Fig. 4b, e) of Ami-*YGL1* lines and WT at different stages (DAT 46, 53 and 60) was measured in field with canopy gas exchange systems CAPTS (Fig. 4c, f) (Song & Zhu, 2018). Results show that the *A*_*c*_ of homozygous lines was lower than WT and the *A*_*c*_ of heterozygous lines was not different from WT at different stages with data at DAT 46 shown in the Fig. 4a and d. Further, whole day cumulative CO_2_ uptake of the whole canopy (*A*_*c,d*_) was calculated from the diurnal *A*_*c*_ curves at different stages and was not significantly different between the heterozygous and WT, while *A*_*c,d*_ of homozygous lines was significantly lower (*P* 0.0015) than that of WT and heterozygous (Fig. 4b, e). Following (Song et al., 2022), we fitted the maximal canopy photosynthesis rate (*P*_*cmax*_) and quantum yield of canopy photosynthesis (*Φ*_*c*_) from the *A*_*c*_-*Q* curve reconstructed based on the *A*_*c*_ and PPFD during a day (Supplementary Fig. S2). *P*_*cmax*_ and *Φ*_*c*_ was not significantly different between the heterozygote and WT and was lower in homozygote than WT (Table 1).

**Table 1.**
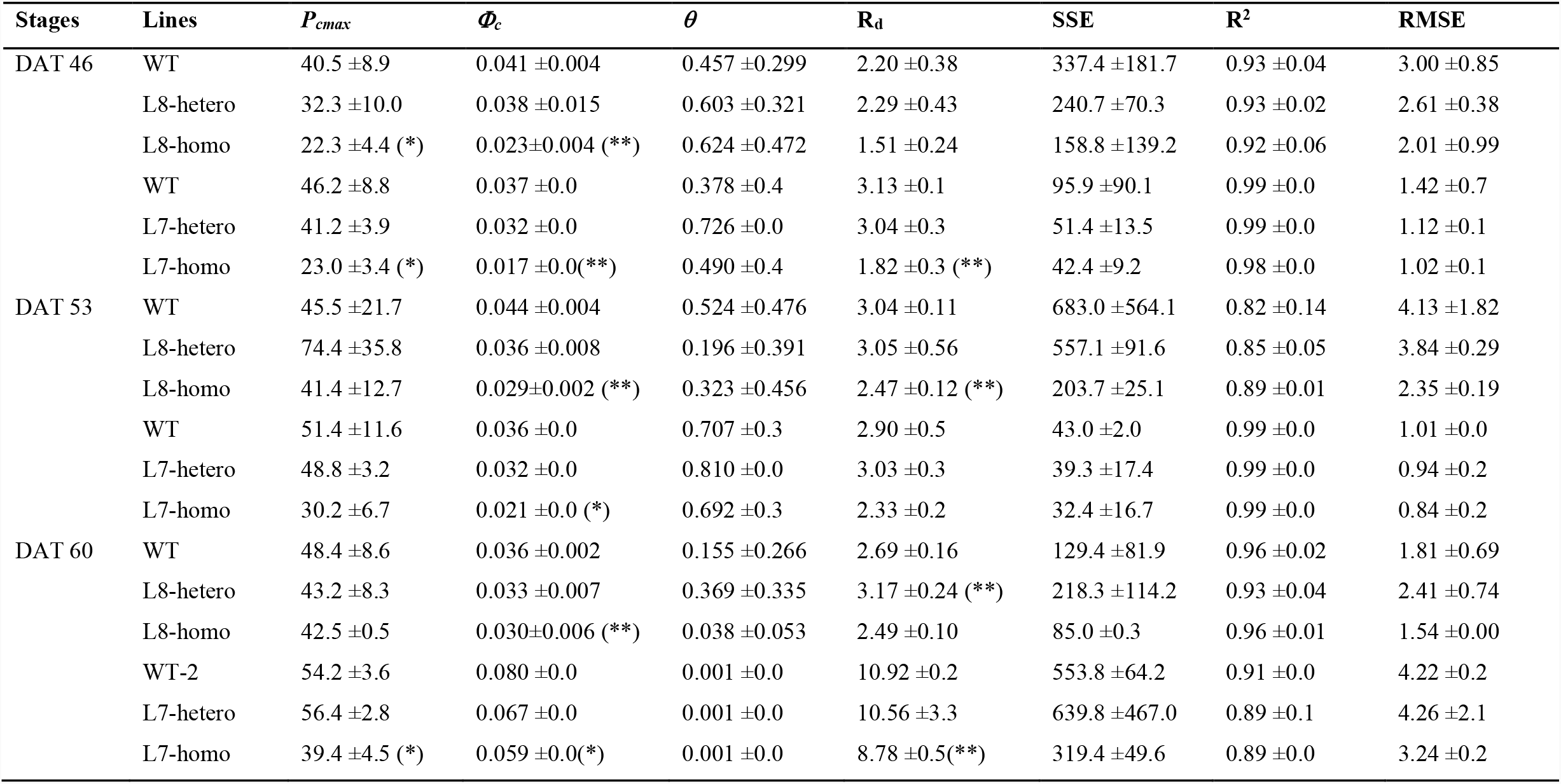
fitting parameters of diurnal canopy photosynthetic CO_2_ uptake rate (A_c_) under different photosynthetic photon flux density (PPFD) for Ami-*YGL1* lines and WT at three stages (DAT 46, 53 and 60). The diurnal data of A_c_ and PPFD measured were used to fit a non-rectangular hyperbola curve. *P*_*cmax*_ is the maximal canopy photosynthesis rate, *Φ*_*c*_ is the initial slope of the curve, *θ* is the convexity of the curve and R_d_ is the canopy respiration rate. SSE, R^2^ and RMSE for the fitting were shown. Data were shown as mean+/−sd with n = 3 biological replicates. Asterisk shows Student’s *t*-test *P*<0.01 (**) and *P*<0.05 (*).

| Stages | Lines | $P_{cmax}$ | $\Phi_c$ | $\theta$ | $R_d$ | SSE | $R^2$ | RMSE |
| --- | --- | --- | --- | --- | --- | --- | --- | --- |
| DAT 46 | WT | 40.5 ±8.9 | 0.041 ±0.004 | 0.457 ±0.299 | 2.20 ±0.38 | 337.4 ±181.7 | 0.93 ±0.04 | 3.00 ±0.85 |
| | L8-hetero | 32.3 ±10.0 | 0.038 ±0.015 | 0.603 ±0.321 | 2.29 ±0.43 | 240.7 ±70.3 | 0.93 ±0.02 | 2.61 ±0.38 |
| | L8-homo | 22.3 ±4.4 (*) | 0.023±0.004 (**) | 0.624 ±0.472 | 1.51 ±0.24 | 158.8 ±139.2 | 0.92 ±0.06 | 2.01 ±0.99 |
| | WT | 46.2 ±8.8 | 0.037 ±0.0 | 0.378 ±0.4 | 3.13 ±0.1 | 95.9 ±90.1 | 0.99 ±0.0 | 1.42 ±0.7 |
| | L7-hetero | 41.2 ±3.9 | 0.032 ±0.0 | 0.726 ±0.0 | 3.04 ±0.3 | 51.4 ±13.5 | 0.99 ±0.0 | 1.12 ±0.1 |
| | L7-homo | 23.0 ±3.4 (*) | 0.017 ±0.0(**) | 0.490 ±0.4 | 1.82 ±0.3 (**) | 42.4 ±9.2 | 0.98 ±0.0 | 1.02 ±0.1 |
| DAT 53 | WT | 45.5 ±21.7 | 0.044 ±0.004 | 0.524 ±0.476 | 3.04 ±0.11 | 683.0 ±564.1 | 0.82 ±0.14 | 4.13 ±1.82 |
| | L8-hetero | 74.4 ±35.8 | 0.036 ±0.008 | 0.196 ±0.391 | 3.05 ±0.56 | 557.1 ±91.6 | 0.85 ±0.05 | 3.84 ±0.29 |
| | L8-homo | 41.4 ±12.7 | 0.029±0.002 (**) | 0.323 ±0.456 | 2.47 ±0.12 (**) | 203.7 ±25.1 | 0.89 ±0.01 | 2.35 ±0.19 |
| | WT | 51.4 ±11.6 | 0.036 ±0.0 | 0.707 ±0.3 | 2.90 ±0.5 | 43.0 ±2.0 | 0.99 ±0.0 | 1.01 ±0.0 |
| | L7-hetero | 48.8 ±3.2 | 0.032 ±0.0 | 0.810 ±0.0 | 3.03 ±0.3 | 39.3 ±17.4 | 0.99 ±0.0 | 0.94 ±0.2 |
| | L7-homo | 30.2 ±6.7 | 0.021 ±0.0 (*) | 0.692 ±0.3 | 2.33 ±0.2 | 32.4 ±16.7 | 0.99 ±0.0 | 0.84 ±0.2 |
| DAT 60 | WT | 48.4 ±8.6 | 0.036 ±0.002 | 0.155 ±0.266 | 2.69 ±0.16 | 129.4 ±81.9 | 0.96 ±0.02 | 1.81 ±0.69 |
| | L8-hetero | 43.2 ±8.3 | 0.033 ±0.007 | 0.369 ±0.335 | 3.17 ±0.24 (**) | 218.3 ±114.2 | 0.93 ±0.04 | 2.41 ±0.74 |
| | L8-homo | 42.5 ±0.5 | 0.030±0.006 (**) | 0.038 ±0.053 | 2.49 ±0.10 | 85.0 ±0.3 | 0.96 ±0.01 | 1.54 ±0.00 |
| | WT-2 | 54.2 ±3.6 | 0.080 ±0.0 | 0.001 ±0.0 | 10.92 ±0.2 | 553.8 ±64.2 | 0.91 ±0.0 | 4.22 ±0.2 |
| | L7-hetero | 56.4 ±2.8 | 0.067 ±0.0 | 0.001 ±0.0 | 10.56 ±3.3 | 639.8 ±467.0 | 0.89 ±0.1 | 4.26 ±2.1 |
| | L7-homo | 39.4 ±4.5 (*) | 0.059 ±0.0(*) | 0.001 ±0.0 | 8.78 ±0.5(**) | 319.4 ±49.6 | 0.89 ±0.0 | 3.24 ±0.2 |

**Figure 4.**
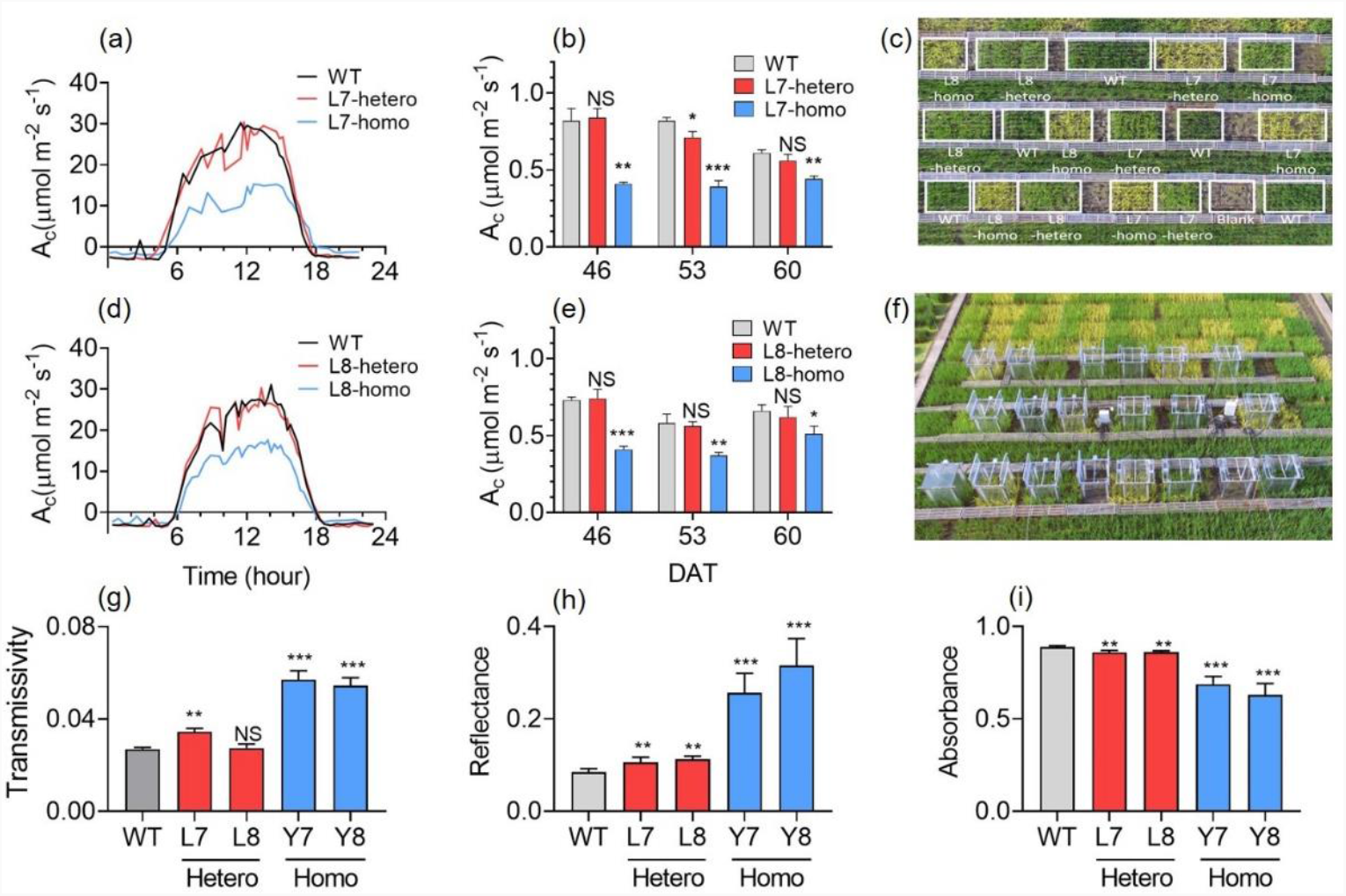
Canopy photosynthesis and canopy light interception. The diurnal canopy photosynthetic CO_2_ uptake rate of Ami-*YGL1* heterozygous lines, homozygous lines and WT **(a, d)**. Daily total canopy CO_2_ uptake (A_c,d_) at different days after transplanting (DAT) **(b, e)**. The A_c,d_ was calculated as the integral of A_c_ during a day (24h). The block design of plots in the field **(e)** and the facilities of canopy gas exchange system CAPTS **(f)** for measuring the A_c_. Canopy transmittance **(g)**, reflectivity **(h)** and absorbance **(i)** of Ami-*YGL1* and WT. Data were shown in mean +/− std (n= 3 plots) and asterisk shows Student’s *t*-test *P*<0.05 (*), *P*<0.01(**), *P*<0.001 (***), NS shows non-significant. All the *t*-tests were performed with WT.

Canopy photosynthesis is mainly determined by leaf photosynthetic efficiency and the total canopy absorbed light. In this study, we measured the canopy absorbance, reflectance and transmittance for the Ami-*YGL1* lines and WT. The Ami-*YGL1* lines showed a higher transmissivity (Fig. 4g), reflectance (Fig. 4h) and less absorbance (Fig. 4i) than WT (*P* < 0.01 for heterozygous, *P*<0.001 for homozygous), except for that the transmittance of line 8 heterozygous line was not significantly different with WT (Student’s *t* test, *P* = 0.59).

### Agronomic traits of Ami-*YGL1* plants in the field

We assessed the effects of reducing chlorophyll content and antenna size on agronomic traits of field grown rice. Plant height, tiller number, panicle number, biomass dry weight, harvest index and grain yield (Fig. 5) were not significantly different between line 7 heterozygote and WT; while tiller number (*P* < 0.01), panicle number (*P* < 0.01), harvest index (*P* < 0.05) and grain yield (*P* < 0.05) of line 8 heterozygote showed statistically significant higher values than WT. Plant height, biomass dry weight, harvest index and grain yield of homozygous lines were significantly lower than WT (*P* < 0.001), showing that the major agronomic traits were greatly affected by large decrease of chlorophyll content and antenna size (Fig. 5).

**Figure 5.**
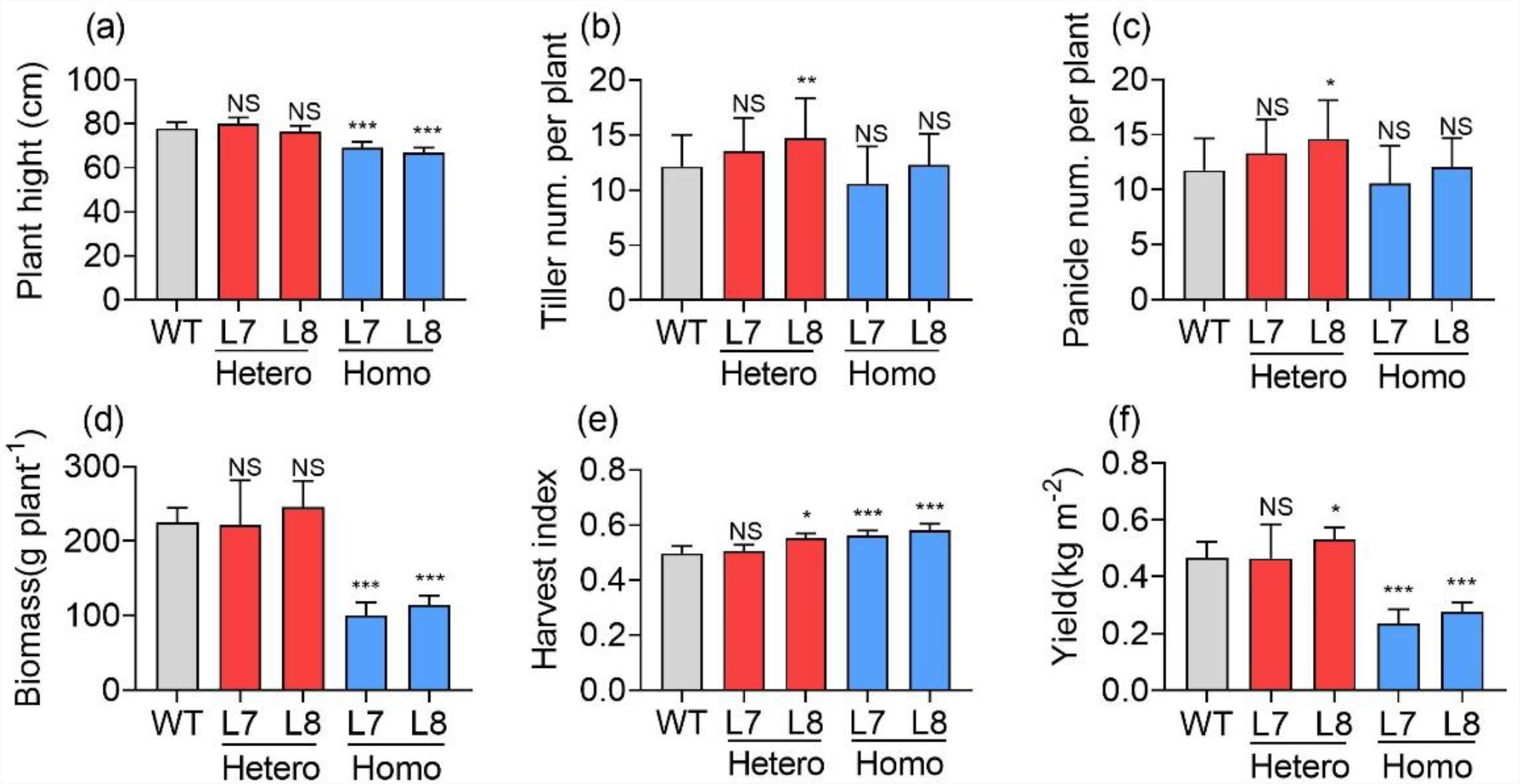
Agronomic traits for Ami-*YGL1* plants and WT. Agronomic traits including plant height **(a)**, tiller number per plant **(b)**, panicle number per plant **(c)**, biomass dry weight **(d)**, harvest index **(e)** and grain yield **(f)** were measured for field grown plants. Asterisk shows Student’s *t*-test *P*<0.05 (*), \*\**P*<0.01, \*\*\**P*<0.001, NS shows non-significant. All the *t*-tests were performed with WT.

### Relationship between leaf photosynthesis, chlorophyll content and antenna size

Our above analysis already shows that the decrease in antenna size can result in different impacts on leaf photosynthesis, and correspondingly agronomic traits (Fig. 4-5). To further study factors controlling these differential responses of changing antenna size on photosynthesis, we chose leaves with varying chlorophyll content from all Ami-*YGL1* transgenic lines including both heterozygous and homozygous plants. The net leaf photosynthesis rate under a PPFD of 1500 μmol m^−2^ s^−1^ (*A*_*1500*_), chlorophyll content (SPAD reading), functional antenna size and F_v_/F_m_ were measured for each leaf at tillering and grain filling stages (Fig. 7a-d). Leaves with F_v_/F_m_ higher than 0.8 were regarded as showing no photoinhibition or photodamage (Piotr Robakowski & Wyka, 2010). We plotted the data at the two stages with F_v_/F_m_>0.8 separately (Fig. 7 e,f) and results show that *A*_*1500*_ did not correlate to chlorophyll content and antenna size when leaves F_v_/F_m_ was higher than 0.8 (Fig. 6e, f), *i*.*e*. when leaves were not under photodamaged state. The relationship between F_v_/F_m_ and *A*_*1500*_ for both two stages was shown in Fig. 7a. When F_v_/F_m_ was lower than 0.8, the *A*_*1500*_ was positively correlated with F_v_/F_m_ (Fig. 7b) and when F_v_/F_m_ was higher than 0.8, *A*_*1500*_ did not correlates with F_v_/F_m_ (Fig. 7c).

**Figure 6.**
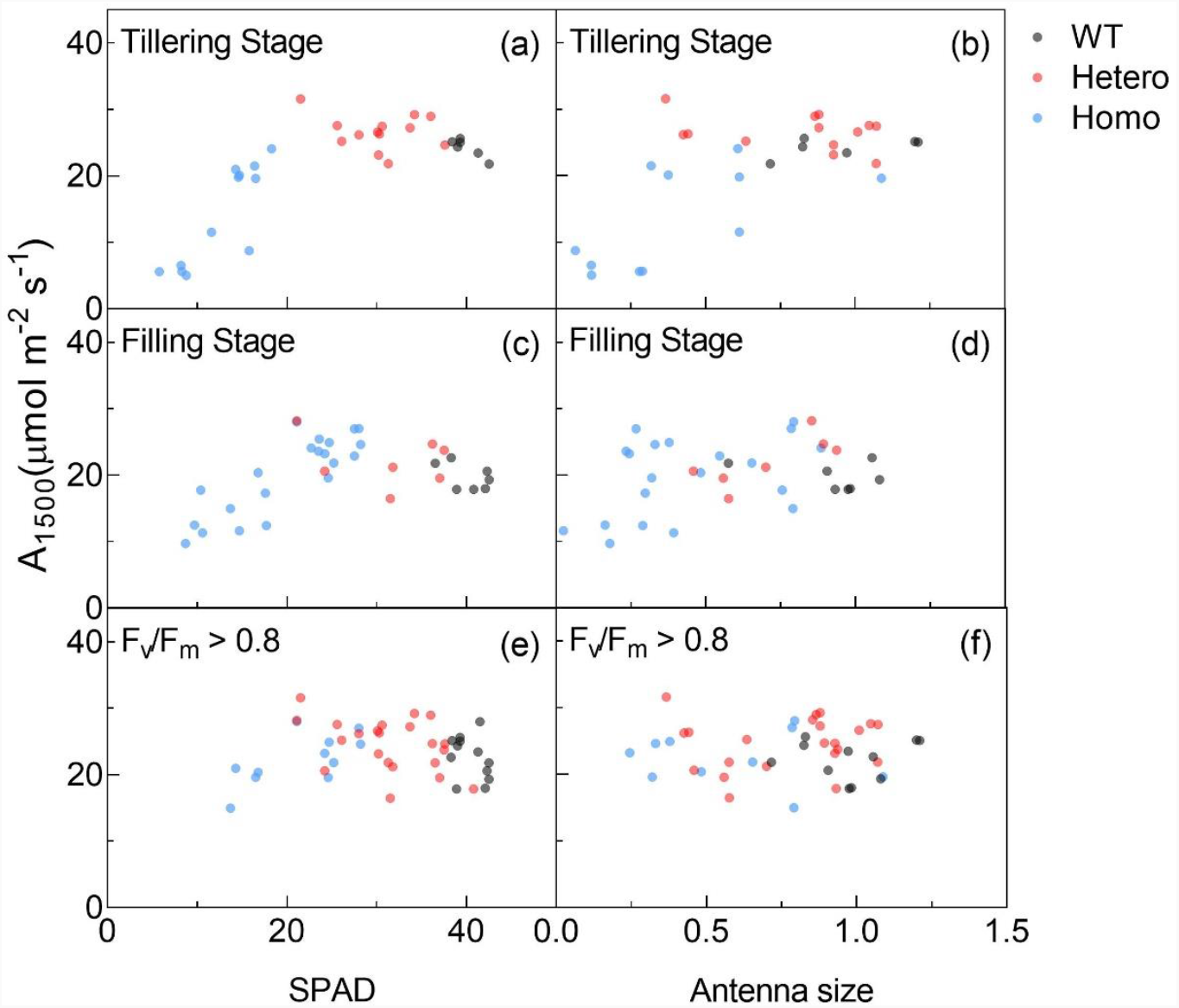
The relationships between leaf photosynthetic rate under saturating PPFD (A_1500_) and chlorophyll content (SPAD) and antenna size. Relationships between A_1500_ and SPAD for Ami-YGL1 lines and WT at the tillering stage **(a)** and the grain filling stage **(c)**. Relationships between A_1500_ and antenna size for Ami-YGL1 lines and WT at the tillering stage **(b)** and the grain filling stage **(d)**. All data at both tillering and grain filling stages with F_v_/F_m_ higher than 0.8 were selected to present the relationship between A_1500_ and SPAD **(e)** and the relationship between A_1500_ and antenna size **(f)** for leaves with Fv/Fm > 0.8, *i*.*e*. under non-photoinhibition state. Black points represent WT, red points represent data from the heterozygous lines and blue points represent data from the homozygous lines.

**Figure 7.**
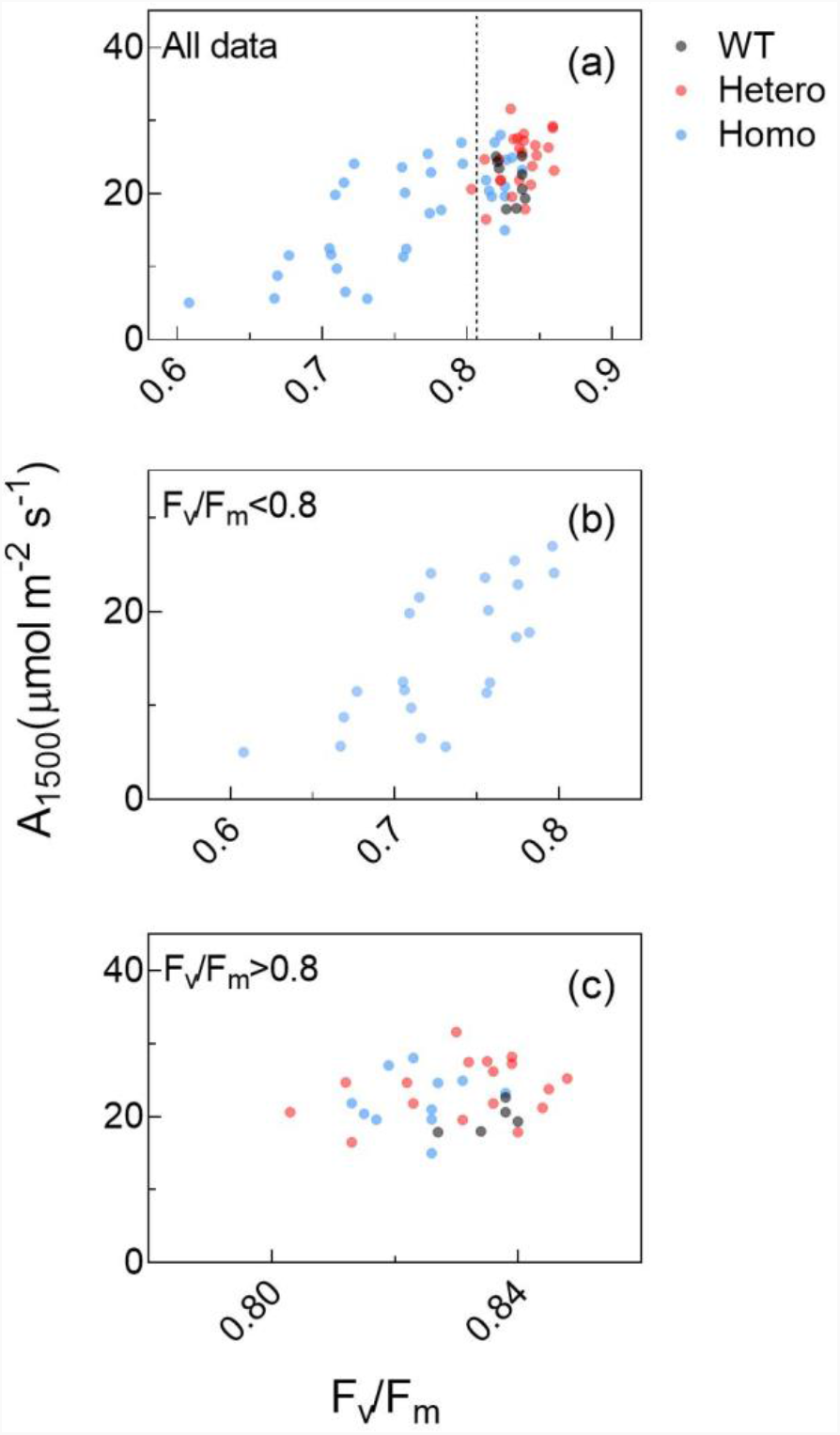
Relationship between A_1500_ and F_v_/F_m_ for all data with F_v_/F_m_ ranging from 0.6 to 0.86 (a), for F_v_/F_m_ lower than 0.8 (b) and for F_v_/F_m_ higher than 0.8 (c).

We found that all the leaves with F_v_/F_m_<0.8 were from homozygous lines, showing that the homozygous lines were prone to photodamage than WT and the heterozygous lines (Fig. 7b). To test this, we treated dark-adapted rice leaves with a PPFD of 1200 μmol m^−2^ s^−1^ for three hours followed by dark recovery for three hours. F_v_/F_m_ was measured during the treatment and recovery phase (Supplementary Fig S3). Result shows that, during high light treatment, the decrease of F_v_/F_m_ for Ami-*YGL1* homozygote line was larger than heterozygote and WT while there was no significantly difference between the heterozygous lines and WT, and the recovery of F_v_/F_m_ in Ami-*YGL1* homozygous lines were lower than that in the heterozygous lines and WT (Supplementary Fig S3).

The above analysis suggests that decreasing antenna size can decrease leaf photosynthetic rate only if the decrease caused decrease in the maximal quantum yield of PSII, *i*.*e*. F_v_/F_m_; when there is no impact on F_v_/F_m_, the antenna size of PSII can be decreased without a negative impact on leaf photosynthesis. We further tested this using current high-yielding cultivars. Specifically, we measured the *A*_*1500*_ and chlorophyll content and antenna size for 6 high-yielding cultivars widely grown in China (Fig. 8). The range of antenna size for these cultivars was between 0.6 to 1.6 and SPAD reading was between 36 to 43 (Fig. 8). First, we found that in general, the *A*_*1500*_ did not show a significant change with changes in either leaf chlorophyll content or antenna size, consistent with the results of Ami-*YGL1* lines when F_v_/F_m_ was higher than 0.8, and *A*_*1500*_ did not correlated with antenna size under this condition (Fig. 8d). Since the leaf chlorophyll content and antenna size of these high yield cultivars were much higher than those in Ami-*YGL1* heterozygous lines, there is still a large space to reduce antenna size in current elite rice cultivars in China without affecting photosynthesis efficiency.

**Figure 8.**
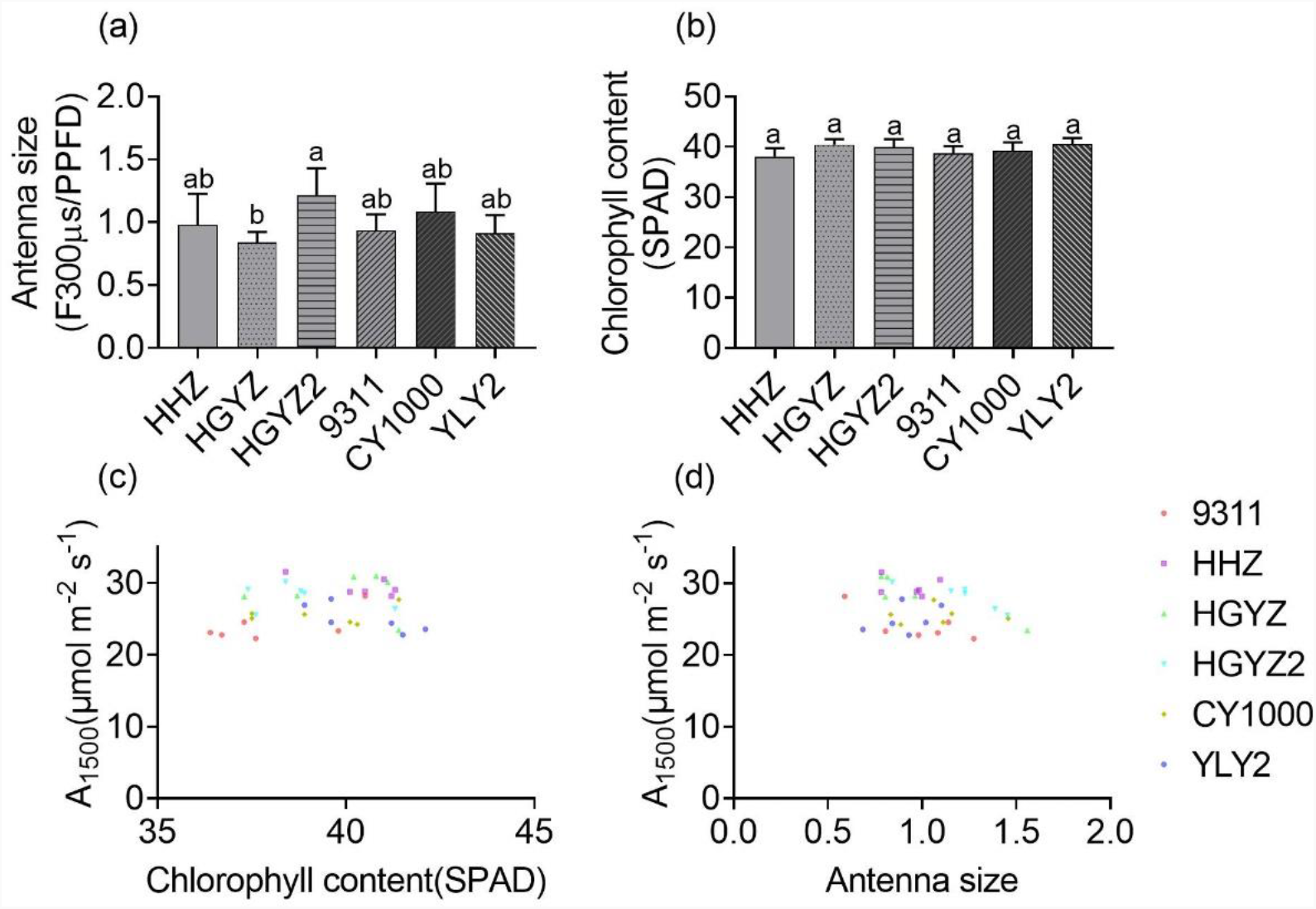
The relationships between leaf photosynthesis (A_1500_) and chlorophyll content (SPAD), A_1500_ and antenna size for high yield rice cultivars. Antenna size and SPAD for high-yielding rice cultivars, *i*.*e*., HHZ, 9311, HGYZ, HGYZ2, CY1000 and YLY2, under a PPFD of 1500 µmol m^−2^ s^−1^**(a, b)**. Relationship between A_1500_ and chlorophyll content **(c)** and relationship between A_1500_ and antenna size **(d)**.

## DISCUSSION

This study shows that the photosystem antenna size can be decreased without negatively influencing the leaf photosynthetic rates as long as the maximal quantum yield of photosystem II can be maintained. Furthermore, we show that there are large variations in leaf chlorophyll contents and antenna sizes, suggesting that there is still large space to manipulate leaf antenna for improved nitrogen use efficiency.

### Antenna size can be decreased effectively through decreasing chlorophyll synthesis

Photosystem antenna system is composed of pigments, mainly Chl, and protein complexes (Masuda et al., 2003, Li et al., 2021). The proportion of Chl assigned to the peripheral antenna protein is affected by leaf Chl content (Petra & Ingo, 2008). In less green leaf mutants, such as Mg-chelatase subunit D mutants, *ygl3* (Tian et al., 2013) and *ygl7* (Deng et al., 2014), the Chl a/b ratio was higher and the content of antenna protein was less compared to WT. Here in our experiments, decreasing the expression of *YGL1* resulted in decreased leaf chlorophyll and antenna sizes (Fig. 2a, c). All these show that decreasing leaf chlorophyll content is a viable strategy to decrease antenna size, while the leaf chlorophyll content can be decreased through manipulation of enzymes involved in the chlorophyll synthesis pathway.

It is worth mentioning here besides modification of genes involved in the chlorophyll synthesis pathway, antenna size can also be reduced by changing the content of antenna protein or even through manipulation of other pigments in LHCII, such as Lutein, Violax-anthin and Neoxanthin (Polle et al., 2001). Many mutants of *lhcbs* indeed showed smaller antenna size compared to their corresponding WTs (Pietrzykowska et al., 2014, Nicol et al., 2019, Chen et al., 2018). However, in other mutants, such as in lhcb3 mutant, Chl *a/b* decreases, suggesting increase in antenna size (Damkjær et al., 2009). Therefore, decreasing the synthesis of chlorophyll can be used as an effective approach to decrease photosystem antenna size.

### Decreasing antenna size might not influence leaf photosynthetic rate as long as the photoprotection capacity is not compromised

Though decreasing antenna size for higher photosynthetic efficiency has been proposed for long, however, the impacts of decreasing antenna size on photosynthesis are still under debate. In most of the cases, the decrease in antenna size leads to improved photosynthesis (Baroli & Melis, 1998, Ort & Melis, 2011, Jin et al., 2016, Kirst et al., 2017, Sakowska et al., 2018, Gu et al., 2017a). However, in some other cases, the decrease of antenna size indeed resulted in no improvement of photosynthesis. For example, in Chlamydomonas, decreasing the antenna size of PSII through knocking out the chlorophyll b synthesis resulted in decreased quantum yield, decreased F_v_/F_m_, and also decreased light saturated rate of photosynthesis when algae are grown under the sole carbon source acetate (Polle et al., 2000). In another mutant which lacks lutein, violaxanthin and neoxanthin, and correspondingly a small PSII antenna size, the photon use efficiency derived from the initial slope of light saturation curve of photosynthesis is similar between WT and the mutant (Polle et al., 2001).

In our experiments in rice, indeed, even in the same genetic background, when there is different degree of decrease in antenna size and leaf chlorophyll content, the leaf photosynthetic rates show differential impacts (Fig. 6). Specifically, under relatively high leaf chlorophyll concentrations or large antenna sizes, decrease in leaf antenna did not decrease or even increase leaf photosynthetic rates (Fig. 6b, d); however, when the leaf chlorophyll content was decreased dramatically, there is a decrease in leaf photosynthetic rates, showing the non-linear responses of leaf photosynthetic rates to changes in leaf chlorophyll concentrations (Fig. 6a, c). Earlier studies show that when the antenna is larger, the excitation energy pressure to the reaction center is higher and more NPQ is conducted to dissipate excess energy as heat to protect the photosystem (Goss & Lepetit, 2015, Ort & Melis, 2011, Spalding et al., 1984). The decreased NPQ in the homozygous line 7 and 8 suggest that in these lines with much decreased antenna size, the photoprotection capacity (NPQ)(Elrad et al., 2002, Li et al., 2009) was indeed greatly compromised (Fig. 3c), which may underlie the observed decrease in Φ_PSII_(Fig. 3d) and also greatly decreased content of proteins involved in the light reactions, including D1, LhcA3, LhcB1, PsbO, PetA, PetC and PsbS (Fig. 2b). To further confirm that the decreased ability to protect or maintain PSII efficiency might have been a factor underlying the decreased photosynthetic rates in the homozygous lines, we plotted the relationship between leaf photosynthesis under high light (A_1500_) and SPAD value or antenna size (Fig. 6). We found that when all leaves were pooled together, there is a non-linear relationship between A_1500_ and SPAD and antenna size (Fig. 6). When only those data points with F_v_/F_m_ higher than 0.8 were used in the plotting, decreasing antenna size did not decrease A_1500_ (Fig 6e, f); when only leaves with F_v_/F_m_ lower than 0.8 were used in the plotting, decreasing antenna size clearly decrease A_1500_ (Fig 7b, c), again showing that the photosynthetic rates did not correlated with antenna size as long as the photosystem II efficiency is well maintained, which most likely as a result of robust capacity of NPQ or photoprotection (Murchie & Lawson, 2013, Krause & Weis, 1991). Besides the decreased chlorophyll content underlying the observed decrease in NPQ (Gu et al., 2017a, Sakuraba et al., 2017), our experiments show that decreased PSBS content might also be another factor (Fig 2b). The content of PSBS protein decreased in the Ami-*YGL1* homozygous lines (Fig. 2), which is different from plants with reduced antenna size reported earlier (Flannery et al., 2021, Demmig-Adams & Adams, 1992, Yu et al., 2021). PSBS is associated with LHCII trimer (Li et al., 2020, Kiss et al., 2008, Rochaix, 2014) and heat dissipation/NPQ (Elrad et al., 2002, Goss & Lepetit, 2015). The decreased PSBS protein contents might have contributed to the decrease the capacity of photoprotection under high light (Fig. S3).

### Canopy with an optimal antenna distribution for greater canopy photosynthetic efficiency

Decreasing antenna size has been shown to be a strategy to improve rice photosynthesis, biomass and productivity(Song et al., 2017, Gu et al., 2017a). Most of these studies evaluated their impacts on leaf photosynthesis and then final agronomic parameters. Here we show that the diurnal canopy photosynthetic CO_2_ uptake rate (A_c,d_) of the heterozygote was not significantly different from the WT but the A_c,d_ of the homozygote was significantly lower than the WT (Fig. 4.a,b,d& e); correspondingly, the plant height, biomass and yield of the heterozygous lines were not significantly different from WT (Fig. 5. a, d and f), while these traits in the homozygous lines were dramatically lower than WT (Fig. 5. a, d and f). Furthermore, the tiller number, panicle number and harvesting index were not decreased in the Ami-*YGL1* transgenic lines (Fig. 5. b, c and e), showing that the manipulation of antenna size by decreasing expression level of *YGL1* did not influence the development of these major canopy architecture parameters. Therefore, this study provides further evidence that decreasing antenna size, as long as it does not compromise the photoprotection capacity, does not lead to decreased canopy photosynthesis. It is worth mentioning here that survey of the SPAD values of elite rice cultivars show that there is a great variation of leaf chlorophyll content in these lines (Fig. 8), suggesting that in modern rice cultivars there is a large scope to decrease leaf chlorophyll content or antenna size for potential higher nitrogen use efficiency without compromising photosynthetic efficiency.

Earlier studies have shown that large antenna is beneficial to plants under low light, *e*.*g*. understory plant species in forest (Evans, 1989), since large antenna can help harvest more light to support photosynthesis under low light (Fig. S4). Plants indeed have evolved to adapt to low light by partitioning more nitrogen to the LHC complex (proteins and chlorophylls) than to the reaction center complex or carbon metabolism enzymes (Emilie et al., 2013). Although the Ami-*YGL1* population had more light distributed to the lower layers in canopy, but the light absorption of these leaves at lower layers is limited due to the decreased antenna size as well (Fig 4 g-i). Therefore, the antenna size of leaves at lower layers of a canopy can be manipulated to maintain a similar or gain even higher antenna size to further improve canopy photosynthesis in the field. Therefore, an ideal canopy might need to have leaves with smaller antenna at the top layers of a canopy and leaves with greater antenna at the lower layers of canopy, *i*.*e*., the smart canopy proposed by Ort *et al*. (2015) can be a feasible strategy to improve crops for higher yield.

### In summary

This study shows that reducing the antenna size is a double-edged sword for canopy photosynthesis. A moderate decrease of photosystem antenna size can optimize light distribution in plant canopy, and hence enhance nitrogen use efficiency; however, when the antenna size is decreased too much, it may compromise the photoprotection capacity and correspondingly results in decreased photosynthetic efficiency and agronomic performance. Furthermore, results from this study provides further evidence for the feasibility of Smart Canopy, in which, leaves at the top layers of a canopy have small antenna size while leaves at lower layers of a canopy have greater antenna size.

## ACKNOWLEDGMENT

This study was funded by National Research and Development Program of Ministry of Science and Technology of China (2019YFA0904600; 2020YFA0907600), Strategic Priority Research Program of the Chinese Academy of Sciences (grant number: XDB27020105) and the general program of National Science Foundation of China (31870214, 31970378) and the Bill and Melinda Gates Foundation (OPP1129902; OPP1172157).

## Abbreviations

Symbol: Description
Chl: Chlorophyll
LHCII: Light-harvesting complex of the photosystem II
CP24: Lhcb6 (24 kDa) protein–pigment complex,a minor LHCII
CP26: Lhcb5 (26 kDa) protein–pigment complex,a minor LHCII
CP29: Lhcb4 (29 kDa) protein–pigment complex,a minor LHCII
D1: D1 protein of the photosystem II core complex
D2: D2 protein of the photosystem II core complex
Cyt b_6_*f*: Cytochrome b_6_*f* complex
PSI: Photosystem I
PSII: Photosystem II
F_0_: Minimum chlorophyll fluorescence
F_m_: Maximum chlorophyll fluorescence
F_t_: Steady-state fluorescence yield
F’_m_: Partial open maximum fluorescence yield
F_v_: Variable fluorescence and it calculated as F_m_-F_o_
F_v_/F_m_: _Maximal_ or potential photochemical efficiency of photosystem II in dark-adapted leaves
Y(II) / ΦPS II: Quantum yields of photosystem II
NPQ: Non-photochemical quenching of chlorophyll fluorescence or excitation energy
ΦNPQ: Quantum yield of light-induced non-photochemical fluorescence quenching
NO: Non-light-induced nonphotochemical fluorescence energy
ΦNO: Quantum yield of non-light-induced non photochemical fluorescence quenching
PPFD: Photosynthetic photon flux density

## Supplemental data

Table S1. Primers used in this study

Table S2. The information of antibodies used in this study

Table S3. The data of the *A-Q* curve at tillering stage.

Table S4. The data and fitted parameters of the *A-Ci* curve at tillering stage.

Figure S1. The measured fluorescence curves for quantification of functional antenna size of PSII.

Figure S2. The fitted light response curve of diurnal canopy photosynthesis.

Figure S3. The maximal photochemical efficiency of PSII measured after dark adaptation of one night (Initial F_v_/F_m_) (a) and the changes of F_v_/F_m_ during high light treatment and subsequent dark recovery (b).

Figure S4. The relationship between leaf photosynthesis rate under low light, chlorophyll content and function antenna size at the tillering stage.

